# Model of multiple synfire chains explains cortical spatio-temporal spike patterns

**DOI:** 10.1101/2022.08.02.502431

**Authors:** Alexander Kleinjohann, David Berling, Alessandra Stella, Tom Tetzlaff, Sonja Grün

## Abstract

It has been postulated that information processing in the brain is based on precise temporal correlation of neural activity across populations of neurons. In a recent study we found spatio-temporal spike patterns in experimental recordings from monkey motor cortex, and here we study if those could be explained by a synfire chain (SFC) like model. The model is composed of groups of neurons connected in feed-forward manner from one group to the next with high convergence and divergence. When activated, e.g., by a current pulse to the first group, spiking activity in the SFC is synchronous within neurons of the same group and propagates from group to group. When a few neurons from different groups are recorded from such an SFC, and the SFC is repeatedly activated, we would find a spatio-temporal spike pattern repeating across trials. Here, we take the statistics of the STPs found in the experimental data from 20 sessions as a reference to compare to a simulated network. Distributions of the data we take into account include 1) the pattern sizes, i.e. the number of neurons involved in the patterns, 2) the number of patterns a single neuron is involved in, 3) the durations of the patterns, and 4) the spatial distances of the patterns across the electrode array used to record the data. For the simulations, we embed SFC(s) in an anatomical model of the respective layer of the motor cortex, defined by its height and the density of the neurons. Model parameters are the length of the SFC, the number of neurons per group, the spatial extent of each neuronal group, and the distance between subsequent groups. Given the size and reach of the Utah array electrodes, we derive the probability of recording neurons from the SFC network. An SFC is considered detected if at least two neurons from two different groups are recorded. We find that depending on the model parameters, an embedded SFC can be detected with high probability, despite the massive subsampling of the cortex by the Utah array. Furthermore, to achieve multiple membership of a neuron in different patterns, we embed multiple SFCs that overlap. The fitting of the model to the pattern data constrains the spatial SFC parameters: the chains have to be broadly distributed in space and contain many neurons per group to match the experimental results.

## 1 Introduction

The synfire chain (SFC) model was proposed by Moshe Abeles as a way to realize a stable and temporally precise propagation of spiking activity in the brain [1] and is based on the complete transmission line, a concept proposed by [2] allowing for a robust transmission of excitation within a randomly connected network of neurons. An SFC consists of multiple groups of neurons. The number of groups is commonly referred to as the chain’s length, and the number of neurons per group as its width. All neurons of a group are connected with high divergence and convergence to the neurons of the subsequent group of the chain (see Fig 1A). If sufficiently many neurons of the first group are stimulated, the elicited spikes arrive at each of the neurons of the next group synchronously and activate these if the dispersion of the spike times is small and the number of spikes is large enough [3]. Thus, such a structure enables a stable propagation of synchronous spikes with millisecond precision. This is further supported by additional studies using a Fokker-Planck approach [4] and allowing for variations in the number of spikes and for temporal jitter in a pulse packet [5]. After the initial proposition by Abeles [1], further theoretical [3,4,6,7] and experimental [8,9] studies showed that in such an architecture, both synchronous spikes and rate profiles propagate through the system with advantages for synchronous propagation [10]. In addition to this robust propagation of activity, SFCs can also be used to gate information which is traveling along another chain, allowing for flexible information routing [11,12]. Interconnected SFCs, each of which is assumed to represent a particular feature, could also serve for binding of information [13]. A visual object can then be represented as a composition of these interconnected SFCs [14,15]. This binding of information may happen in multiple cortical areas in parallel, for which a representation of the information in spatially propagating structures is advantageous [16]. In this context, it has been shown that many SFCs can be embedded in a single cortical network with stable dynamics [13,17,18], and synchronization of SFCs has been proposed as a mechanism for composing information into a more abstract single representation [13,19,20].

**Fig 1.**
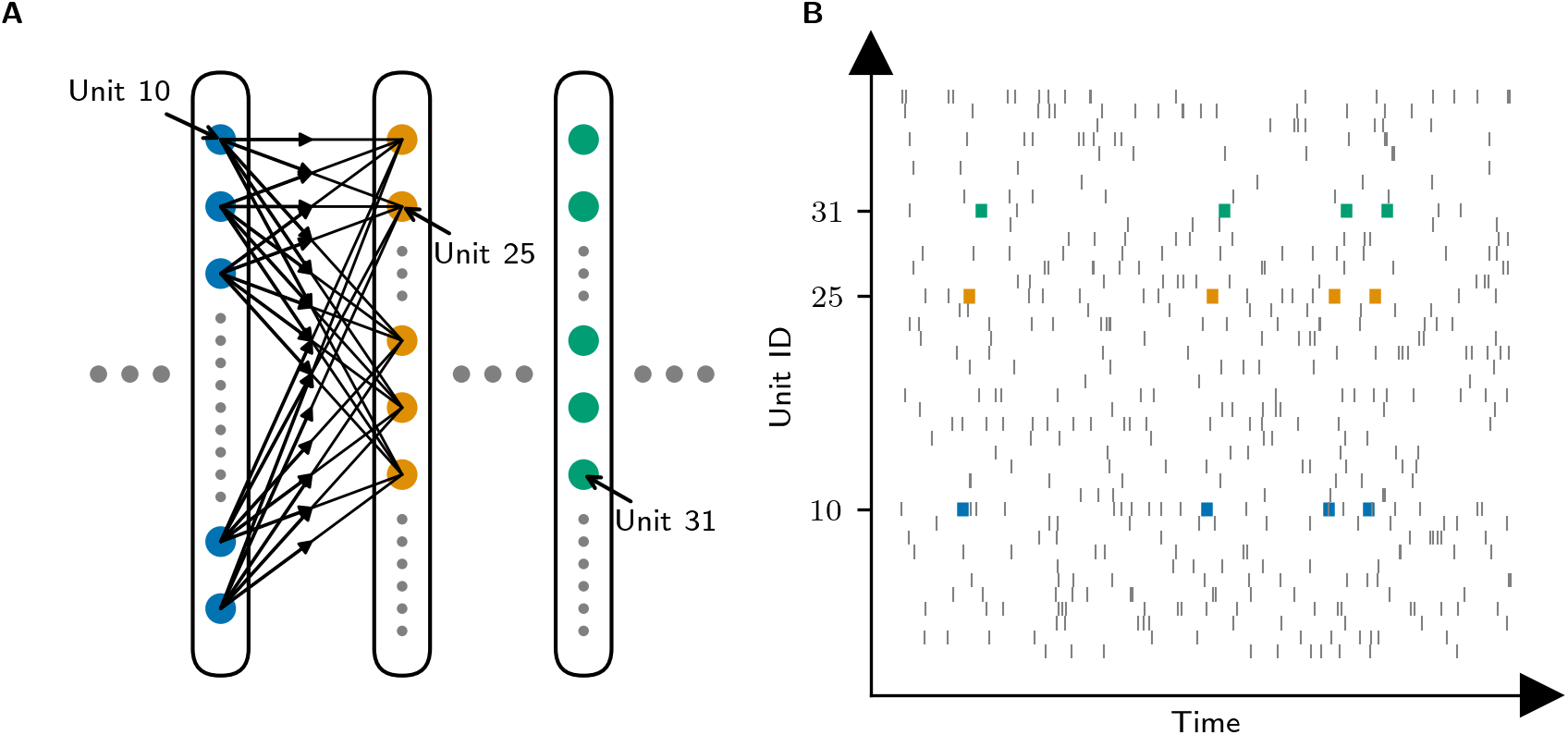
Synfire chain (SFC) model and resulting spatio-temporal patterns (STPs). (A) SFC model. Neurons are connected in groups (colors), here with all-to-all connections from each neuron of the previous group to the neurons of the next group. Once the first group is stimulated with a strong and narrow pulse, synchronous spikes are sent as input to each neuron of the next group, which activates the next group synchronously. (B) The sketch shows neurons potentially recorded from the chain (colored spikes; see arrows in A to unit 10, 25 and 31). Such neurons then spike whenever the SFC is activated, following the order of the group activation, and with a temporal delay corresponding to the signal propagation, thereby forming STPs (blue, yellow, green, 3 times). Grey ticks represent spikes recorded from neurons not in the SFC.

In recent years, a large number of theoretical studies proposed alternative neuronal network models predicting the emergence of precise firing patterns by biophysical mechanisms that are different from those of the classical SFC model of [1]. Similar to the original SFC model, they often rely on successively activated groups of synchronously firing neurons [21–26]. In this work, we therefore refer to these models as SFC-type models, even though the mechanisms triggering synchronous firing in groups of neurons are different from the classical SFC model, the connectivity structure is not always strictly feed-forward, and the group sizes can be substantially smaller (of the order of 10) if nonlinear dendritic integration (dendritic action potentials) is accounted for [21–23,26]. Other models predict the emergence of robust precise firing patterns without synchronous firing, but with specific temporal delays, such as the “braids” model [14], or the polychronization networks [27].

Temporal and spatio-temporal patterns are observed across several species and brain areas, and at different temporal resolutions ranging from milliseconds to seconds [16,28–39]. In a recent study we found spatio-temporal spike patterns (STPs) in experimental data recorded from primary motor cortex (M1) of two monkeys [40]. Here, we investigate if these patterns can be explained by SFCs, given the recording setup with a 10×10 Utah array. Our assumption is that an STP results from recordings of at least two different neurons that take part in the SFC (as sketched in Fig 1B). We conceptually embed SFC(s) into a cortical volume corresponding to the region recorded from by the Utah array: an area of 4 × 4 mm^2^, layer 2/3 and its respective neuron density. Additionally, the sensitivity of the individual electrodes of the Utah array are considered for estimating the neurons recorded from in the model.

The distributions of the number of neurons involved in an STP, the temporal extents of the STPs, the spatial cortical distance of neurons involved in STPs, and the involvement of a single neuron in different STPs are the references for comparison. As shown in the modeling results (Section 3.1.2), a single SFC is likely to be detected for a large range of model parameters. However, to account for the STP characteristics found in the experimental data, the model has to include multiple SFCs within the assumed piece of cortex. Given a conservative estimate of how many chains exist in the recorded volume, the detection of SFCs in form of STPs is very likely despite the massive subsampling by the Utah array (Section 3.1.3). Additionally, the fact that individual neurons occur in different patterns requires that these SFCs overlap and include the same neurons (Section 2.3). The final results of the simulations (Section 3.2.4) correspond well to the experimental data, which leads to the conclusion that STPs are well explained by multiple SFCs.

## 2 Methods

### 2.1 Experiment and data

We are interested in understanding whether STPs detected in experimental data may be explained by the SFC model. Thus, we examine spatio-temporal spike patterns from numerous recording sessions, and consider the recording device, electrode depth and brain area of the experiment for the assumptions of our model to test if the detected patterns can be explained by the SFC model.

The experiment consists in a delayed reach-to-grasp task performed by two Macaque monkeys *(macaca mulatta*,) [41]. Recordings are obtained through a chronically implanted a 10 × 10 multi electrode Utah array (Blackrock Microsystems, Salt Lake City, UT, USA, www.blackrockmicro.com) in the pre-/motor cortex (M1), along the central sulcus. Each electrode has a length of 1.5 mm, and a distance of 400 μm to the neighboring electrodes (ordered in a grid). Given the depth of the electrode tip, the neurons detected are most likely deep in layer 2/3 [42–45], where the neuronal density in primary motor cortex of macaques is between 21,400 mm^-3^ and 40,000 mm^-3^ [46]. The density in layer 2/3 is 18 % larger than the average density across all layers [42,43,45]. In a separate study [40], we analyzed 20 experimental sessions (10 per monkey), recorded in different days over the time span of months. For monkey N, the recordings were performed in 2014, from June to July; whereas, for monkey L, recordings started in October 2010 and finished in February 2011. Each session has a duration of approximately 15 minutes and consists of around 120 successful trials. The data from each session were spike sorted independently using the Plexon spike sorter (Plexon Inc, Dallas, Texas, USA, version 3.3), retaining only the single unit activities (SUAs) with a signal-to-noise ratio ≥ 2.5, and with an average firing rate of < 70 Hz across the trials. Artifacts consisting of hyper-synchronous spikes at sampling resolution occurring across electrodes were assumed to be cross-talk artifacts and were therefore detected and removed in a preprocessing step [31]. Over all sessions, the average number of SUAs per session is 107, however, it differs across monkeys: higher for monkey N (mean =143 ± 14.82 units), and lower for monkey L (mean =70.5 ± 13.93 units). The average number of SUAs per electrode is 1.1 averaged over the two monkeys.

The two monkeys, L and N, were trained to self-initiate trials by pressing a start button. After a waiting period of 400 ms, a visual cue (yellow LED) was shown to the monkey (waiting signal). The monkey was instructed to wait again for 400 ms, until it was presented to another visual cue, lit for 300 ms (from CUE-ON to CUE-OFF). The task consisted in reaching and grasping an object with the indicated grip type and force level. The grip can be either be a precision grip or a side grip. The CUE-ON signal contained the grip information. The monkey then waited for 1, 000 ms, eventually receiving the GO-SIGNAL, which contained the information on the amount of force to exert to pull the object towards itself. The behavioral conditions were selected randomly for each trial. The requested grip and force had to be maintained for 500 ms, and if it was performed correctly, the monkey received a reward. Further details on the experimental setup can be found in [41, 47] and analysis results on the same data are presented in [31, 48].

### 2.2 Analysis approach for spatio-temporal spike patterns in the experimental data

We used the SPADE [49–51] method to detect STPs in the parallel spike trains of these recordings. The method consists of three successive steps. The first is the detection of all putative patterns in the data at a certain temporal resolution, during which the putative patterns are stored along with the information on when and how often they occur. This is achieved by applying a Frequent Itemset Mining algorithm [52, 53]. The second step is the statistical evaluation of the significance of the patterns detected in the first step, under the null-hypothesis of mutual independence of spike trains given their firing rate (co-)modulations [54]. The third step is a conditional test performed on all significant patterns, in order to remove patterns arising from the overlap of true pattern spikes and chance spikes. SPADE outputs all patterns labeled as significant, i.e. STPs, by the statistical tests. Each pattern is returned together with the neuron ids involved in the patterns, the lags between spikes, the times of the occurrences, and the p-value. Details of the method and on its implementation can be found in [51]. The temporal resolution of the detected patterns is given as an input to the SPADE method, and is here fixed to *b* = 5 ms. Moreover, we fix the maximal duration of STPs to 60 ms.

Each recording session contains four trial types (corresponding to the combinations of the two force types and two grip types). For the pattern analysis, we segmented the trials of about 4 s into six 500 ms-long behavioral epochs in each single trial; *start, cue, early delay, late delay, movement* and *reward.* The respective epochs of the same trials of the same session are concatenated and analyzed as an independent data set, and thus in total yield (4 trial types × 6 epochs) data sets per session (see [31] for details on the data segmentation into trial epochs). The data sets of each session are analyzed separately, since the probability to record the same neuron(s) across sessions is unlikely [55].

### 2.3 Reference statistics of STPs

We analyzed the data recorded in the delayed reaching-and-grasping task for STPs using SPADE. We detected 119 patterns in total (monkey N: 61, monkey L: 58) in 20 sessions. The statistics of the results pooled over all sessions are displayed in Fig 2. STPs in different epochs are not identical, but specific to the behavior. However, for this study we pool across trial types, epochs, and sessions. Typically we find around 6 patterns per session (mean = 5.95 ± 3.26). Patterns are formed by spikes emitted by different neurons, and contain spikes of 2-6 neurons (mean = 2.9 ± 0.93 neurons), but the majority of patterns consist of two and three spikes (Fig 2A). Another statistic is the number of different STPs a single neuron (within a single session) is involved in (Fig 2B). Most STP neurons participate in only 1 pattern, but many also in 2-3 patterns, and in a few cases, a single neuron can partake in up to 9 STPs [40, 56].

**Fig 2.**
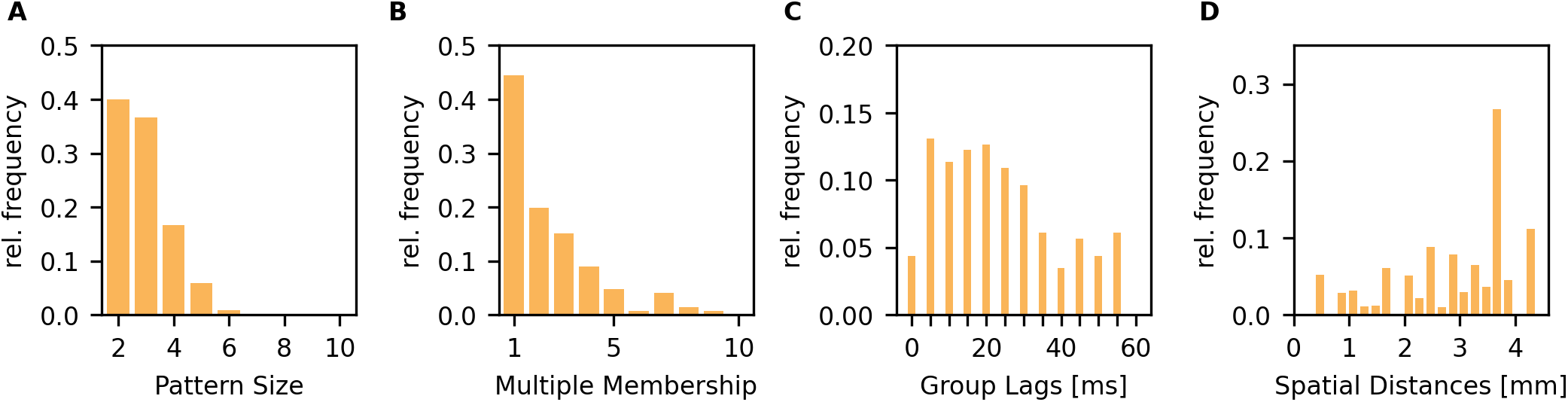
Pattern statistics in the reach-to-grasp experiment of two monkeys, of ten sessions each. (A) Histogram of the number of neurons involved in a STP (pattern size). (B) Histogram of the number of STPs a single neuron appears in (multiple membership). (C) Histogram of time lags between neighboring pattern spikes, i.e. only the time difference between subsequent pattern spikes are taken into account. The temporal resolution of the histogram coincides with the resolution of the SPADE analysis (5ms). The maximum time lag corresponds to the maximal pattern duration (here 60ms). (D) Histogram of the euclidean distances between pattern spikes. Only subsequent pattern spikes are taken into account. The entries for each spatial distance are weighted by the occurrences of the respective distance on the Utah array electrode grid. All histograms are normalized such that the sum of all entries is 1.

Regarding the temporal delays between the spikes in STPs, we observe that the whole range of allowed delays (0 to 60 ms) is covered for both monkeys (Fig 2C). We do not see a particular tendency towards preferred delays or oscillatory activity. However, the results show a slight preference for temporal delays around 10 to 30 ms. We also calculate the euclidean spatial distances between STP neurons of the ones forming successive spikes. The position of the respective neuron is given by the electrode the neuron is recorded from. The distribution of these distances is shown in Fig 2D). This distribution is normalized by the number of distances that exist on the array. The resulting distribution is rather flat, with a peak around 3, 600 μm, however, we tend to ignore that peak since the larger the distances, the fewer number of samples are available which makes the statistics less reliable.

### 2.4 Model for spatial embedding of SFCs in a cortical network

We aim to assess the detectability of SFCs in the experimental setting presented in Section 2.1. Since the recording was performed with a 10 × 10 electrode Utah array recording with 96 active electrodes arranged at 400 μm distance, we only look at the cortical volume below the area of the electrode array (4 mm×4 mm) for this study. Moreover, we only consider layer 2/3 by setting the height of the simulation volume to its thickness: *h*_SFC Volume_ = 1.5 mm (see [42–45]). The volume in which neurons can be detected by the electrodes of the array is confined to spheres with a radius corresponding to the sensitive range (see Section 2.5). The simulation volume and its height *h*_SFC Volume_ = 1.5 mm corresponds to the volume in which SFCs are embedded in our model. Since the sensitive range of the electrodes is a separate parameter, increasing the volume height effectively increases the subsampling of the neurons in the simulation volume by the Utah array. We choose a default value of *ρ* = 35,000 mm^-3^ for the neuron density in our simulation, but we will investigate the impact of varying it in the following sections. In our simulation, we first fill the volume under the Utah array with neurons according to the neuron density (Fig 3A). Then, we simulate a single SFC realization by positioning a sequence of SFC groups in the space of the cortical area. In particular, the SFC embedding procedure starts by drawing the position of the first group center within the borders of the electrode array from a two-dimensional uniform distribution covering the area of the array. Around the first group center, we select *w* neurons inside a cylinder with radius *r*_group_ and height *h*_SFC Volume_ (Fig 3B). The parameter *w* controls the number of neurons per group, and *r*_group_ represents the radius of the cylinder in which all *w* neurons of a single group reside. For the subsequent group, the center position is drawn from a two-dimensional, symmetrical Gaussian distribution which is centered on the previous group center (Fig 3C). The standard deviation *σ*_group distance_ of this Gaussian distribution determines the average spatial distance between groups (〈*d*〉):

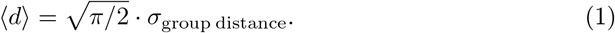

**Fig 3.**
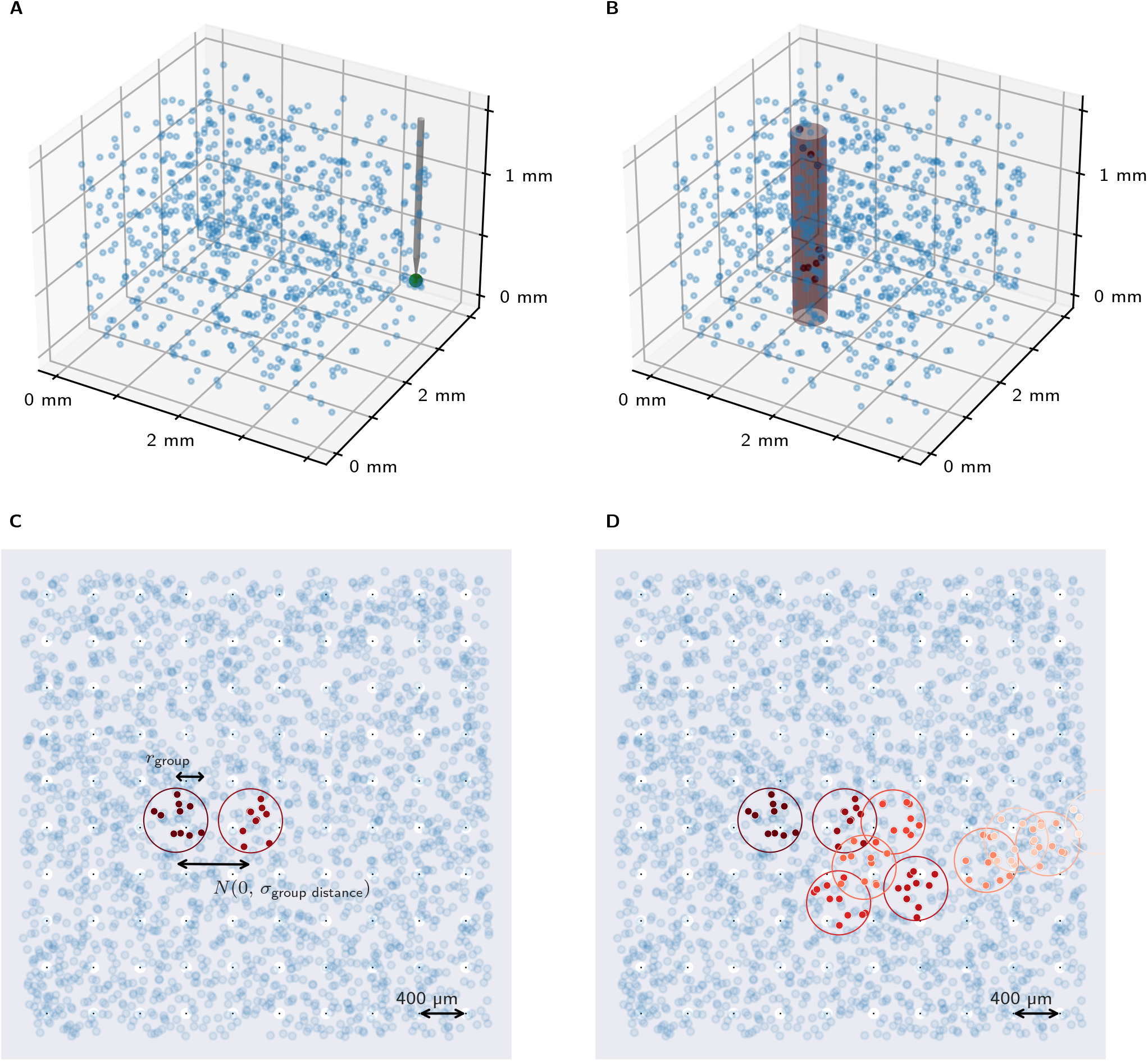
Visualization of the spatial embedding procedure of an SFC. (A) The simulated volume is homogeneously filled with neurons according to the local neuron density (here the density is reduced to 10 mm^-3^ to unclutter the plot). Its size is given by the area covered by the Utah array along the *x* and *y* axes, and by the thickness of the cortical layer 2/3 (1, 500 μm) in which the electrode tips are placed in the *z* axis. To illustrate the relation of the shown spatial scales to the one of an electrode and its sensitive range one electrode (at the corner of the Utah array) is shown in gray on the right side of the plot and its sensitive range is illustrated as a green sphere at its tip. (B) The center of the first synfire group is randomly placed inside the volume. A group is confined by a circle with radius *r*_grcup_ in the horizontal plane, and by the thickness of the cortical layer along the third dimension, resulting in a cylinder (not to be confused with a micro-column). *w* neurons inside the cylinder are randomly assigned to the group. For illustration purposes we show only groups of *w* =10 neurons here. (C) Top down view on the cube showing everything projected onto an *x-y* plane. The Utah electrodes and their sensitive ranges are illustrated across the whole area by black dots surrounded by white circles, respectively. The center of the second SFC is drawn from a two-dimensional Gaussian centered on the previous group center, with a standard deviation *σ*_grcup distance_. The neurons of the second group are assigned analogously to the first group. (D) This procedure is repeated for *l* groups, resulting in one SFC being embedded in the simulated volume.

The next group of neurons is selected inside the cylinder around the center position analogously to the first group. This procedure is repeated to distribute *l* groups iteratively, where *l* represents the length of the SFC, i.e. the total number of groups in the chain (Fig 3D). We allow groups of a chain to be partially or fully outside of the simulation volume (e.g. the final group of the SFC shown in Fig 3D), since there is no plausible reason for SFCs to be constrained to the volume below an electrode array. However, these neurons are not available to be ‘recorded’ from.

To reduce computational cost and account for participation of one neuron in multiple SFCs we always embed multiple SFCs per simulation. In order to achieve that, we repeat all of the above steps *n*_chains_ times to embed *n*_chains_ SFCs in the volume. A neuron can be part of multiple groups and multiple chains. This setup allows us to study multiple trials if we consider single chains one after another.

### 2.5 Neuron isolation model

Since an electrode only records neurons whose soma is close to the electrode tip, we need to consider their distance to the electrode [57]. We make the simplifying assumption that for isolating the activity of a single neuron, its cell body needs to lie within a certain radius around the electrode tip, which we refer to as the electrode’s sensitive range (*r*_sens_):

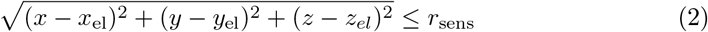

Here, *x, y* and *z* are the spatial coordinates of the neuron and *x*_el_, *y*_e_ and *z*_el_ are the coordinates of the electrode tip (the tip is where the actual recording takes place). We set the depth of the electrode tips to *r*_sens_. This is arbitrary, but *z*_el_ does not have any impact on the results since the *z* coordinates do not affect the assignment of neurons to SFC groups (see Section 2.4). Thus, a neuron can be recorded if it lies within a sphere with a radius equal to the sensitive radius of the electrodes (*r*_sens_). The experimental data show that on average 1.1 single units are recorded from a single Utah array electrode, estimated from all 20 sessions (see Section 2.1). Thus, we set the isolation probability of a neuron located in the sensitive sphere around an electrode tip with radius *r*_sens_ accordingly:

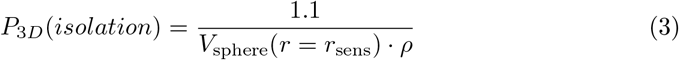

with the neuron density *ρ* in the vicinity of the electrode and the number of single units per electrode in the numerator. Note that this means that independently of the SFC parameters, the constant average number of detected neurons per electrode is always 1.1. This matches the number of isolated single units, which might seem low at first glance. However, to be selected, they also have to be isolated by the spike sorting software, which requires that neurons have to fire often enough, have a high enough signal to noise ratio and have a spike waveform that is clearly distinguishable. This number is thus to be expected to be quite low, up to 10- to 100-fold lower than expected from the neuron density [58]. Assuming a sensitive radius of *r*_sens_ = 50 μm [57] and a neuron density of *ρ* = 35,000 mm^-3^ [46], there are 18 neurons in the sensitive volume of each electrode of which we can isolate 1.1 per electrode, matching the 10- to 100-fold subsampling observed by Shoham et al. (2006) [58]. See [47] for more detailed information on the applied spike sorting procedure.

#### Sensitive Range of Utah Electrodes

In order to estimate the sensitive range of the Utah array electrodes, we have to extrapolate from measurements in a different setting due to the unavailability of directly comparable data. [57] estimated the sensitive range of extracellular electrodes to be 50 μm by comparing simultaneous extracellular and intracellular recordings. The electrodes they used have a significantly higher impedance than Utah array electrodes (400 kΩ to 600 kΩ vs 50 kΩ, cf. [59]), which results in a smaller sensitive range and a higher signal-to-noise ratio, but they were also recording in rat hippocampus which has a ten-fold higher neuron density than macaque M1 [60,61] which we are considering in our experimental setting (cf.Section 2.1). Since these differences have both positive and negative contributions to the sensitive range, we assume a range of 50 μm for most analyses but investigate the effect of smaller (30 μm) and larger ranges (70 μm) on the detectability of SFCs to quantify the associated uncertainty. For illustration of the size relationships we refer to Fig 3A, which shows a sketch of an electrode of the Utah array including its sensitive range (green sphere).

### 2.6 SFC detectability measure

With our model for spatial embedding of synfire chains (cf. Section 2.4) we simulate the positions of the neurons of the chain. Assuming a 10×10 Utah array and the isolation model (cf. Section 2.5) we determine whether these or some of these neurons are recorded from. Repeating this experiment (embedding and measuring) with the same parameters yields probability distributions for recording different numbers of neurons given these parameters.

Next, we would like to quantify the ability to detect an SFC in such a setting. For that, we assume the dynamics in such an embedded SFC as determined by [3] for the case of stable propagation. Diesmann et al. [3] assumed an all-to-all connectivity from one group to the next and found that at least 80 neurons are required per group to get a stable propagation. If the chain is activated by a strong current pulse in parallel to all neurons of the first group, this first group then exhibits synchronous firing such that all neurons of the next group are also activated synchronously at some delay (corresponding to the time delay from one group to the next). That way, a packet of synchronous activity propagates through the chain. For the detection of an active SFC, we require that at least two neurons from two different groups are recorded, since only spike patterns with delays show propagation of activity along the chain. Thus, we define the SFC detectability as the probability to detect neurons from two or more different groups, cf. Fig 4. The detected neurons correspond to an STP where the time difference between spikes of the different neurons are assumed to be given by the number of SFC groups between the two neurons multiplied by the time to propagate from one group to the next, here assumed as 5 ms. With these assumptions, we get one STP per SFC.

**Fig 4.**
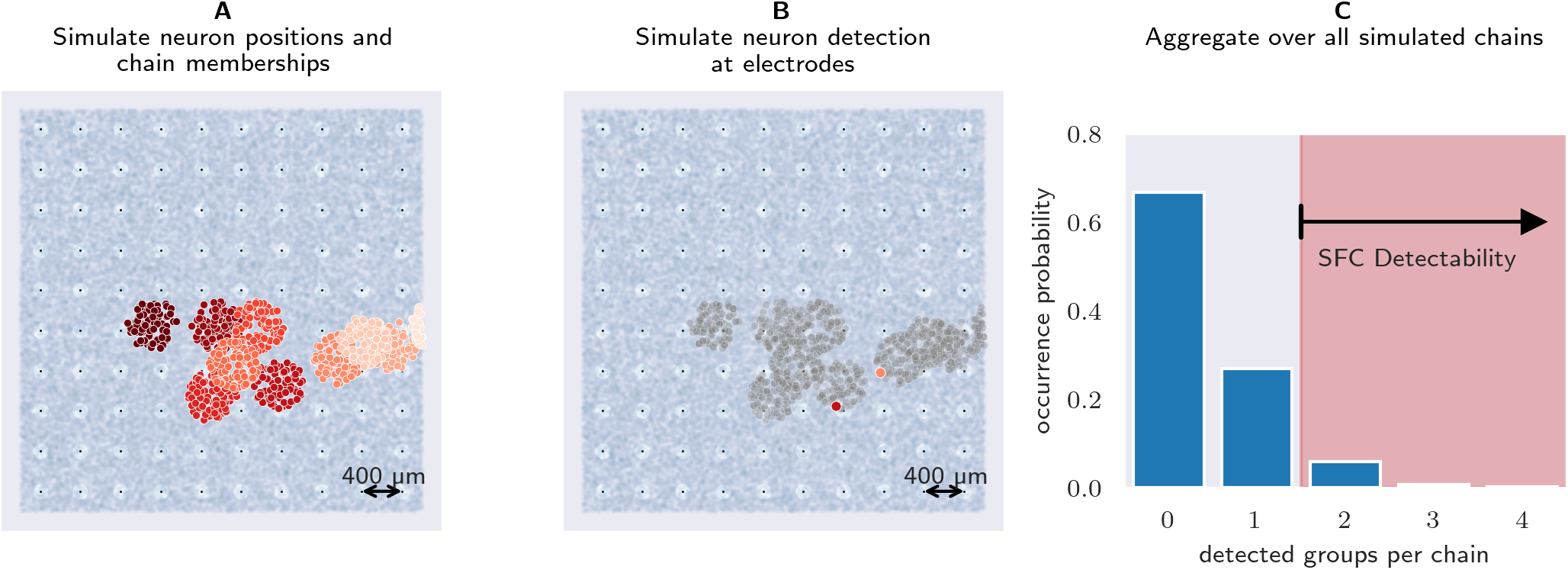
SFC detectability measure. (A) Neuron positions on the 4 × 4 mm^2^ array in color-coded groups for an exemplary SFC sample. The color shows the SFC group, lighter colors correspond to later groups. The black dots surrounded by a white circle indicate the recording electrodes, and the radius of the circle indicates the sensitivity range of the electrode. (B) Same type of plot as in A, but only the neurons recorded by the electrodes are marked in color. (C) Histogram of the probability to record a certain number of neurons per embedded SFC, pooled across 1,000 embedding experiments. The cumulative sum from 2 detected neurons per SFC and more yields the probability for SFC detection, here 0.065.

## 3 Results

In this section, we first present the results of the STP analysis of experimental data, and show the corresponding statistics. Since we assume that the subsampling of an SFC may lead to such spike patterns, we aim to verify that SFCs can be detected given the experimental Utah array recording setup described in Section 2.1 and our spike pattern analysis (Section 2.3). Thus, we come back to our model for spatial embedding of SFCs (see Section 2.4), and verify the influence of model parameters and recording constraints on the probability of observing a single SFC (SFC detectability). Within the parameter regions with good SFC detectability, we compare the predicted spike pattern statistics obtained by multiple simulations of our model to the statistics obtained experimentally. This yields the best-fit parameter set in which our model can explain the spike patterns in experimental data assuming that the underlying network utilizes SFCs for information propagation.

### 3.1 SFC detectability

Our next question is if one embedded SFC is detectable in 10×10 Utah array recordings. In that context we first aim to answer how the different SFC parameters and recording constraints influence the SFC detectability.

#### 3.1.1 Influence of single model parameters on SFC detectability

We start by fixing all but one parameter and investigate the effect of the varied parameter on the SFC detectability. Therefore, we have to decide on reasonable default values for all parameters. We assume that the temporal delay for the assumed propagation of the SFC activity from one group to the next is *b* = 5 ms. In the experimental data we found STPs with a maximum temporal delay of 60 ms, which was the maximal allowed duration of the patterns in the analysis. This corresponds to a maximal SFC length of *l* = 12, since *l · b* =12 · 5 ms = 60 ms (see Section 2.3). For the SFC width we choose for a start an intermediate value of *w* = 1, 300 which is larger than the minimal value *w* = 100 for stable, robust propagation of synchronous spiking along the SFC [3] but far away from the maximal value of the number of neurons that fit into the smallest assumed group disk *r*_group_ = 300 μm we consider. The maximal value would be 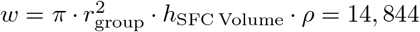 neurons per group. Getting close to this maximum would also mean that most neurons within *r*_group_ would be inside the same group of the same SFC, since no other neurons would be available anymore in the volume and thus would not allow for any additional embedding flexibility.

The spatial parameters *r*_group_ and *σ*_group distance_ are limited by two aspects: very small values would result in SFC groups that fit in between the electrodes’ sensitive ranges and in an SFC that does not move spatially, respectively. On the other hand, very large values would result in parts of a group or parts of the chain residing outside the volume below the area of the Utah array. Therefore, we fix both parameters to an intermediate value of *r*_group_ = *σ*_group distance_ = 900 μm. For the sensitive range of the electrodes we choose *r*_sens_ = 50 μm, but we expect this not to have an effect as discussed in the appendix (S1 Appendix: The influence of the sensitive range of the electrodes).

The measures required and computed in this section only require single embedded chains, however, we embed *n*_chains_ = 10,000 SFCs within one simulation (of 20) since that is computationally much more numerically efficient than performing 200, 000 simulations for the same sample size. We have verified that separate simulations with just a single chain provide similar results (not shown). With these default values for our parameters, we can now investigate the effects of all parameters (one by one) on the SFC detectability.

The SFC length *l* determines the number of groups in an SFC. Increasing it trivially increases the amount of neurons in the SFC and thus the detectability increases (Fig 5A). For high values of *l* this effect flattens out since longer chains have an increased probability of leaving the simulation volume, and thus reduce the number of neurons that can be recorded.

**Fig 5.**
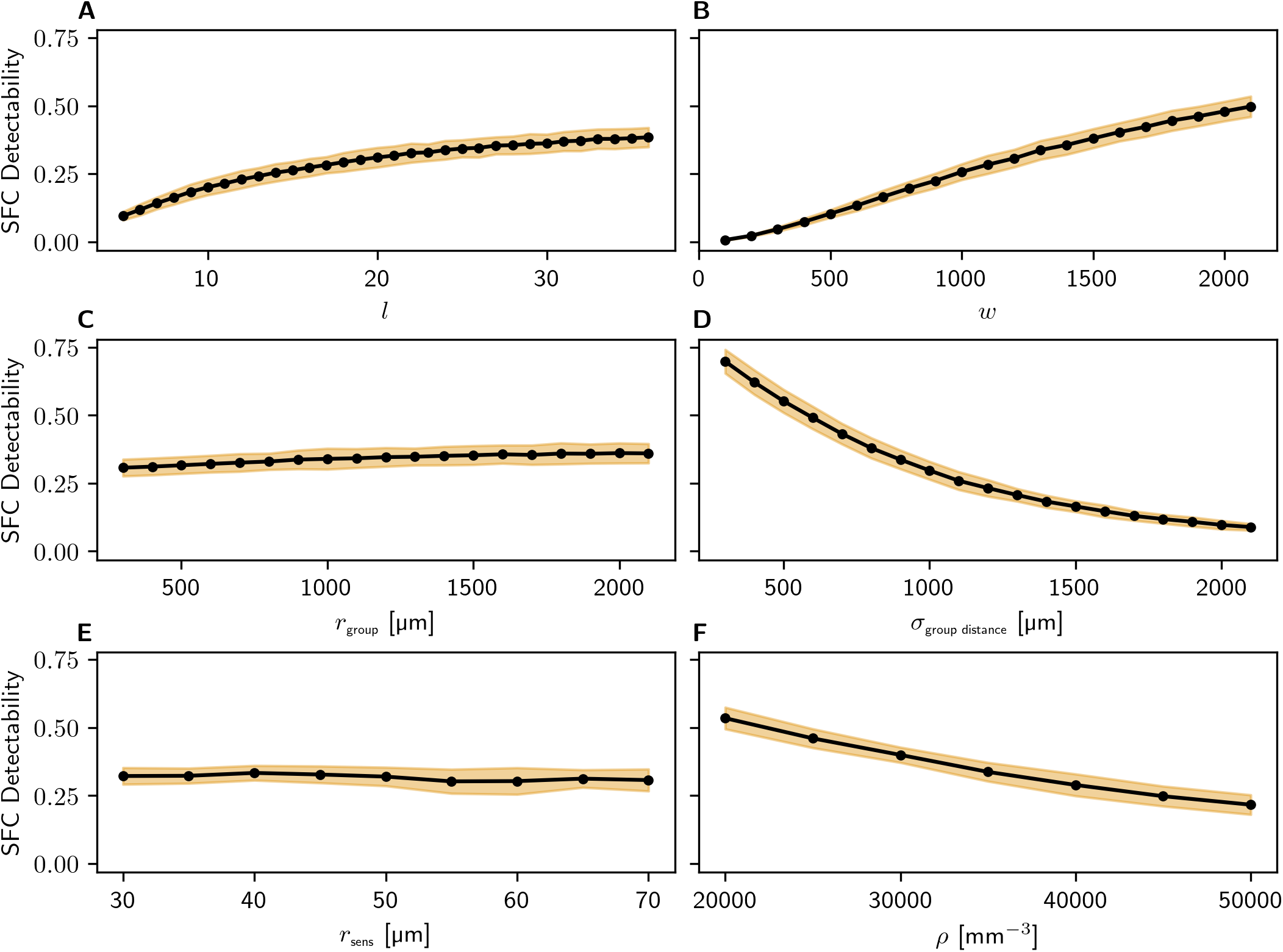
Effects of single parameters on SFC detectability. SFC Detectability as a function of: (A) SFC length *l* in steps of 1; (B) SFC group width *w* in steps of 100; (C) Spatial SFC group size *r*_grcup_ in steps of 100 μm; (D) Spatial SFC inter-group distance *σ*_group distance_ in steps of 100 μm; (E) Sensitive range of the electrodes *r*_sens_ in steps of 5 μm; (F) Neuron density in steps of 5, 000 mm^-3^. Orange bands represent standard deviations over 200, 000 evaluated embeddings.

The SFC width *w* determines the number of neurons in the SFC groups and increasing *w* also leads to an increase of the detectability, however with a different slope. That curve does not flatten out, since for example, a higher *w* for a given *l* just adds more neurons to the existing groups and thus further increases the detectability (Fig 5B).

The spatial SFC parameters *r*_grcup_ and *σ*_group distance_ have quite different effects on the detectability. The spatial SFC group size *r*_grcup_ hardly affects the detectability (Fig 5C). The increased area covered by an SFC group is counteracted by the decreasing density of SFC neurons inside that area, such that the detectability only increases very slowly for larger radii. That slow increase is however consistently monotonous. In contrast to that, increasing the spatial SFC inter-group distance *σ*_group distance_ decreases the detectability (Fig 5D), since a higher inter-group distance results in a higher chance for the SFC to leave the simulation volume. The supplementary figure (S2 Figure: Impact of SFCs leaving the simulation volume) supports this argument; if the boundary conditions for the recorded area/volume were periodically continued there would be no decay of SFC detectability.

The sensitive range of the electrodes *r*_sens_ has no effect on the SFC detectability in our model (Fig 5E). This seems rather counter-intuitive, but the sensitive range is implicitly included in the number of SUAs that can be recorded from an electrode (Eq 3). This parameter has to be fixed in our model since we have to select the number of neurons that can be recorded by an electrode. In order to do this we have to set a maximal distance *r_sens_* from the electrode tip. The exact value we choose does not matter since it eventually cancels out as discussed in the appendix (S1 Appendix: The influence of the sensitive range of the electrodes).

We vary the neuron density between *ρ* = 20,000 — 50, 000 mm^-3^ to account for the measurement uncertainty and the variance between measurements in different studies [42,43,45,46]. The neuron density has a straight-forward effect on the SFC detectability. Increasing the neuron density results in more neurons in the simulation volume but decreases the ratio of neurons within one SFC to total neurons. This results in a decreasing detectability (Fig 5F).

#### 3.1.2 SFC detectability across the entire combined parameter space

In the previous section, we derived how single parameters affect the SFC detectability. For this, we fixed all but one parameter and investigated the effect of the varied parameter. In order to check for possible co-dependencies, we will now vary all parameters at once and calculate the SFC detectability for parameter combinations across our parameter space.

In Fig 6 we represent the results of 720 parameter configurations varying all parameters. We run the simulated model 20 times for each configuration with 10, 000 chains per simulation, and we fix the non-SFC parameters to the worst case scenario of high neuron density *ρ* = 50, 000 mm^-3^ and a high thickness of layer 2/3 of *h*_SFC Voiume_ = 1, 600 μm. The parameters varied simultaneously are: the SFC length *l,* the SFC group width *w* in the *y* axis; the SFC group disk *r*_group_ and the s.d. of the spatial displacement of SFC groups *σ*_group distance_ along the x-axis. The SFC detectability is represented by color (note the color bar): light/dark blue color corresponds to a low/high probability of SFC detection (probability of detecting neurons from two or more different SFC groups).

**Fig 6.**
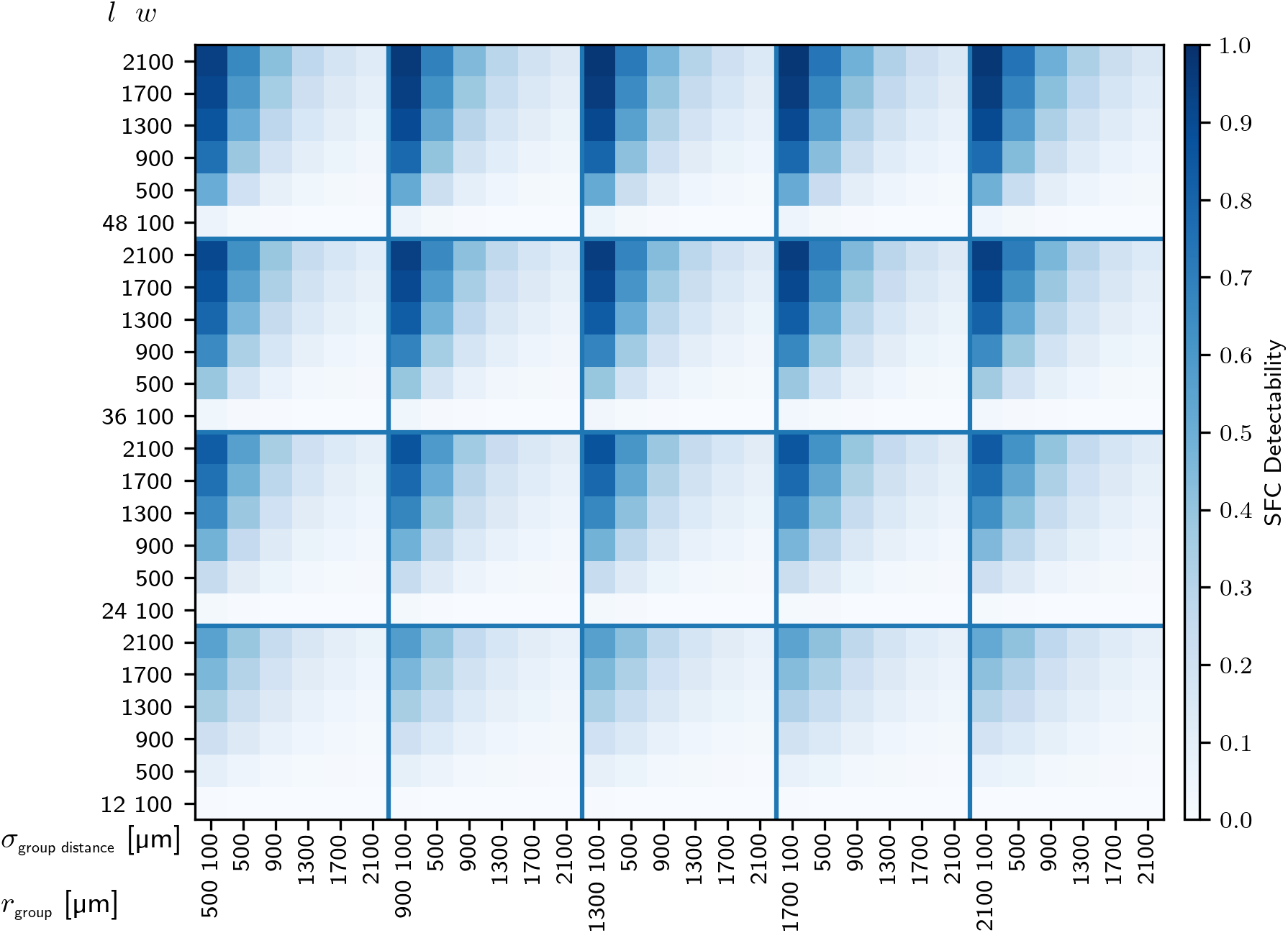
Detectability of a single SFC within an SFC volume height of 1, 500 μm. The color indicates SFC detectability (see color bar) i.e. the probability to detect 2 or more neurons in different groups of a single SFC. Length *l* and width *w* of the SFC are displayed on a combined y-axis, with *l* varying from one quadrant to the next, and *w* varies along each quadrant. The SFC detectability characteristics of varying the spatial distribution determined through the group disk radius *r*_group_ (changing from one quadrant to the next) and the s.d. of the group distance *σ*_group distance_ (changing within each quadrant) are shown along the x-axis. Each entry corresponds to *N* = 20 runs of the model with *n*_chains_ = 10,000 SFCs embedded, i.e. 200,000 evaluations of a single SFC.

Increasing the group width *w* (along the y-axis within each quadrant), the SFC detectability increases. If in addition the SFC length *l* is increased (across the quadrants along the y-axis) the SFC detectability increases even more. Intuitively, the more neurons belong to an SFC, the more likely it is that two neurons are detected from different groups. Increasing the SFC group displacement *σ*_group distance_ (along the x-axis within the quadrants) lowers the SFC detectability for a given fixed group disk radius *r*_group_, whereas a change of the group disk radius *r*_group_ (across the quadrants along the x-axis) does not lead to big differences.

In summary, the SFC detectability is highest for high *l* and *w* and low group distances *σ*_group distance_. The SFC detectability reaches a maximum of 0.975 for *l* = 48, *w* = 2,100, *σ*_group distance_ = 100 μm and *r*_group_ = [1, 300, 1, 700, 2,100] μm. This means that for such values the detectability of an SFC is very likely, and we should observe a spatio-temporal pattern of at least two neurons involved. These parameter configurations with large groups are actually feasible and lead to good agreement with experimental data as will be seen in the following sections.

#### 3.1.3 SFC detectability depending on the number of present SFCs

In the previous section we estimated the detectability of a single SFC which lies in the cortical volume where the Utah array is recording from. However, assuming that there is more than one SFC in the recorded cortical volume, the probability of detecting an SFC should be larger. As explained in Section 2.6 each SFC results in one STP if at least two participating neurons are recorded and cannot result in multiple STPs since they would simply be merged into a larger STP in the spike pattern detection process. Thus also the probability to detect spatio-temporal spike patterns should be larger if multiple SFCs are embedded and assumed to be activated. We can estimate the probability of recording at least one STP by considering an anatomically realistic estimation of the number of SFCs under the Utah array and combining this number of present SFCs with the detection probability of one single chain.

To obtain a conservative estimate of how many SFCs we expect in the cortical volume which is accessible with the Utah array we assume the following: only excitatory neurons form SFCs and each neuron takes part in only one SFC. An SFC consists of *l* groups with *w* excitatory neurons per group. Hence, the number of neurons in each SFC is

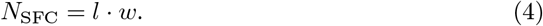

The volume in which we can measure SFCs is given by the area of the Utah array in the horizontal plane (*A*_array_) times the SFC volume height (*h*_SFC volume_) along the vertical direction. Based on the neuron density and the evidence showing that about 80% of all neurons are excitatory [62—65], we can estimate the number of neurons inside the volume we record from (*N*_SFC volume_):

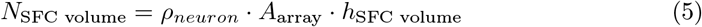

For an SFC volume height of 1, 500 μm and an intermediate neuron density of *ρ_neuron_* = 35,000 mm^-3^, we get a total of 840,000 neurons and *N*_SFC volume_ = 672,000 excitatory neurons in the simulation volume. Dividing this neuron count by the number of neurons per SFC (*N*_SFC_) yields the maximum possible number of SFCs inside the volume:

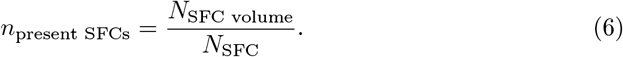

Thus for our parameter ranges that would result in 7 to 560 chains. The probability of detecting at least one SFC when multiple SFCs are present is binomially distributed:

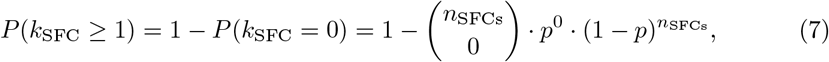

where *n*_SFCs_ is the number of SFCs in the cortical volume and *p* is the probability to detect a single SFC (the SFC detectability). The required number of SFCs in the cortical volume to detect at least one SFC with a probability equal to or greater than *α* is obtained by plugging *P*(*k*_SFC_ ≥ 1) = *α* into Eq 7 and solving for *n*_SFCs_:

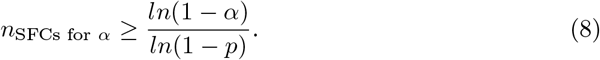

For an example parameter set with *l* = 12 groups, *w* = 100, *r*_group_ = 900 μm, *σ*_group_ = 900 μm which is constrained to the extent of layer 2/3 (SFC volume height of *h*_SFC volume_ = 1, 500 μm), the probability of detecting a single chain is *p*(SFC detected) = 0.0708 (simulation result from Section 3.1.2) and the number of SFCs is *n*_present SFCs_ = 560 (obtained by plugging in the example parameters in Eq 4, Eq 5 and Eq 6). By plugging this into Eq 8 we get minimal number of SFCs needed to detect at least one SFC with a 99% probability: *n*_SFCs for *α*_ =99% ≥ 63.

Fig 7 shows the probability of detecting at least one SFC in simulated Utah array recordings. While the detection probability for an individual SFC can range from 0.0002 to 0.975 as we saw in the previous section, the probability of detecting at least one SFC stays above 95% (*α* ≤ 0.05) for the majority of parameter configurations. This is because our conservative estimate of the number of present SFCs is very high, which compensates for the low detectability of an individual chain.

**Fig 7.**
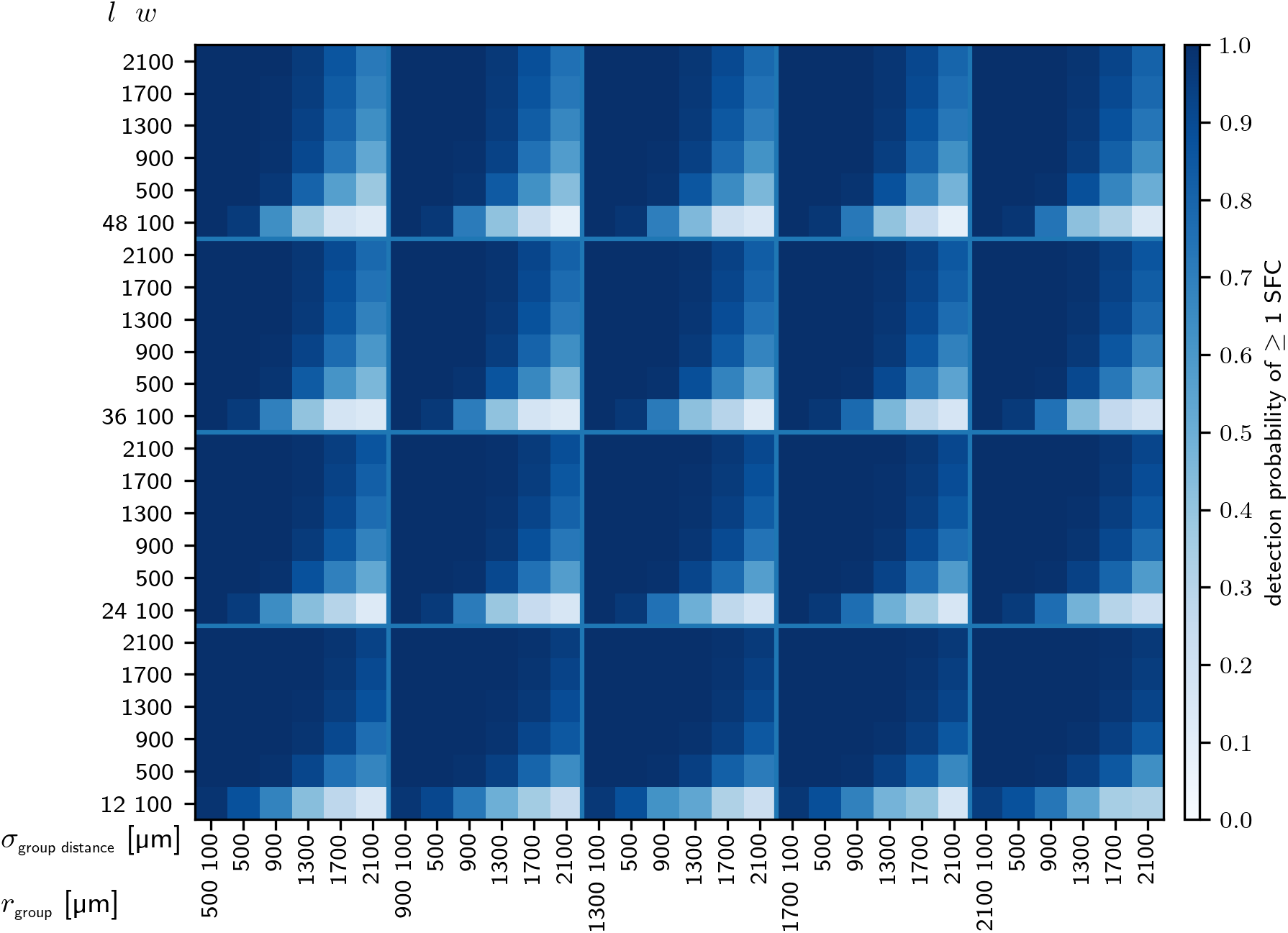
Detectability of ≥ 1 SFC in Utah array recordings in layer 2/3 of macaque M1. Similar visualization to Fig 4. The color denotes the probability to detect at least one SFC for the respective parameter set. Each entry corresponds to N=20 runs of the simulated model.

### 3.2 Comparison of spatio-temporal patterns in SFC model and experiment

The detectability measure we introduced in Section 2.6 was designed to give a binary yes-or-no answer to the question whether the SFC can be detected. To this end, we required that two neurons from different groups of the SFC are recorded. Beyond that, more insights can be gained from considering the exact number of neurons that are expected to be recorded from an SFC and their distribution over the groups of the SFC. This reveals the statistics of the spike patterns that are expected to be found as a result of the SFC. With our model we are able to determine the pattern sizes, which are given by the number of recorded neurons per SFC. In addition, we can turn this around and check for each detected neuron in how many detected chains it appears. Here we allow neurons to participate in multiple chains to match the multiple participation of single neurons in patterns observed in the experiment (see Section 2.3). The assumption in the previous section that every excitatory neuron is part of exactly one chain was just used to get a conservative estimate of the number of present SFCs.

Each detected chain results in one spike pattern. Within such a pattern we can determine the distribution of spatial distances between neurons emitting subsequent pattern spikes. By plugging in an assumed synaptic delay between groups of 5 ms [66, 67], the times between pattern spikes can be calculated as well. This yields all the metrics presented in Section 2.3 for the patterns detected in experimental data.

#### 3.2.1 Euclidean distance as a measure of histogram similarity

In order to compare the spike pattern statistics predicted by our model to the statistics observed in the experiment, we need a way to quantify their similarity. We will compare the distributions of the four measures presented in Fig 11 for the experimental data: pattern sizes, multiple participation of neurons in patterns, time delays between pattern spikes, and spatial distance between pattern neurons. Commonly, two-sample tests are used for this kind of comparison [68]. However, discrete distributions, which we want to compare here, complicate the tests [69]. An additional caveat is that two-sample test p-values do not penalize a small sample count, i.e. comparing a histogram of 1,000 samples to one with just one sample can still lead to a small p-value if the single sample fits the distribution of the 1, 000 samples. This could happen if for example only one pattern is detected in the model. In such a case, we would like to see this reflected in the similarity measure. Furthermore, it is non-trivial to combine the p-values for the four measures in a final combined measure [70]. That is why we decided to use Euclidean distances of the histogram bins, which correspond to Euclidean distances of all possible discrete values of the distributions. This distance is calculated as

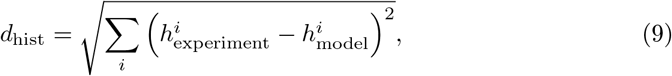

where 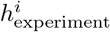 and 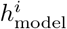 are the values of the ith entry, i.e. the height of the *i*th histogram bar, of the experimental and modeled histogram, respectively. Since all histograms are normalized such that the sum of all of their entries is 1, the total Euclidean distance between two such histograms is always constrained to 0 ≤ *d*_hist_ ≤ 2. This ensures that we can get a combined measure to which all histograms contribute similarly (independently of their number of bins) by simply summing all distances.

We will use the Euclidean distances *d*_pattern size_, *d*_multiple membership_, *d*_group lags_ and dspatial distance and the total distance dtotalin the following.

#### 3.2.2 Influence of single model parameters on spike pattern statistics

Now we look at the effects of single parameters on the Euclidean distance between the distributions resulting from the simulated model and the distribution of pattern statistics observed in experimental data (Section 2.3). In Fig 8, we represent the distances defined in the preceding section in function of five parameters: SFC length *l*, SFC width *w*, SFC group radius *r*_group_ separately, and s.d. of the Gaussian distribution sampling the displacements between SFC groups *σ*_group distance_.

**Fig 8.**
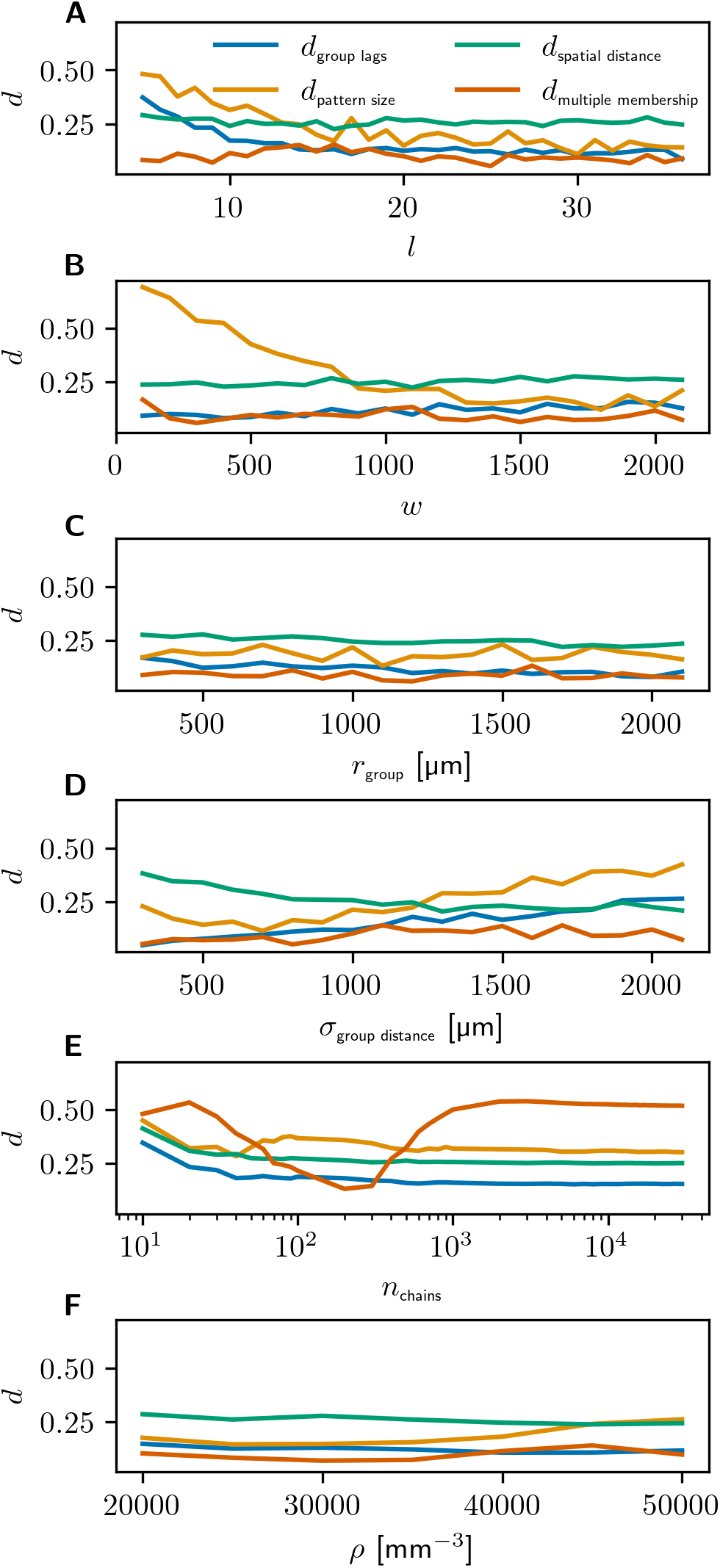
Effects of single model parameters on the euclidean distance between the model results and the pattern statistics. Each distance is calculated between the distribution observed in the experimental data and the one observed in the model. All individual histogram distances (*d*_pattern size_ in orange, *d*_multiple membership_ in red *d*_group lags_ in blue and *d*_spatial distance_ in green) are shown as a function of (A) the number of groups *l*, (B) the group size *w*, (C) the SFC group radius *r*_group_, (D) the SFC group distance *σ*_group distance_, (E) the number of chains *n*_chains_, and (F) the neuron density *ρ*.

We use the same default parameter values as in Section 3.1.1. First, we vary the SFC length and measure how the histogram distance changes as a function of it (Fig 8A). If we assume the transmission delay from one group to the next to be 5 ms, we need *l* ≥ 12 groups in order to generate the longest patterns that can be detected with the SPADE window size of 60 ms. In fact, *d*_grcup lags_ is highest for the lowest values of *l*, decreases for higher lengths and flattens out around *l* = 15. Increasing the SFC length beyond this minimum value results in more patterns and longer delays. This effect saturates at some point and few long chains become indistinguishable from many shorter ones due to the limited SPADE window size. No patterns longer than 60 ms can be detected, but going beyond *l* = 12 does still improve the fit since it increases the chance of observing patterns with delays close to the maximum of 60 ms which are actually observed in the experimental data. This is because there are more possibilities to observe two neurons that are 12 groups apart in a chain of length *l* ≥ 12 than in a chain of length *l* =12. We also observe that *d*_pattern size_ decreases with increasing *l* (orange line in Fig 8A). For very short chains not enough large patterns are found to match the experimental data. Making the chains longer alleviates this issue. This effect also saturates around *l* = 12 since due to the 60 ms duration limit we cannot indefinitely make an STP larger by making the corresponding SFC longer. The other histograms do not show strong effects when varying *l* since they only depend on the total number of chains and the spatial spread of SFCs as we will see in the following paragraphs.

We have already seen that increasing the SFC width results in an increase of detection probability of an SFC. In Fig 8B, we observe that when increasing the SFC width *w* there is an increase in the probability of observing larger patterns. *d*_patternsize_ between the model and the distribution of the SPADE analysis decreases until *w* = 2,000. The other three distances remain constant when *w* is changed.

The SFC group radius *r*_grcup_ determines the spread in space of an SFC group: all neurons of a group are horizontally confined to a circle with radius *r*_grcup_. Results obtained in Fig 2C show that small group radii lead to low similarity between the model and the experimental results. There is a small continuous decrease in both *d*_grcup lags_ and *d*_spatial distance_ with increasing *r*_grcup_. This is probably due to the fact that a small group radius coincides to an increase of synchronous patterns of neurons recorded at the same electrode, since it becomes more likely to detect multiple neurons of a group at the same electrode. However, synchronous patterns and short spatial distances between pattern spikes are not prominently observed in the experimental data (Fig 2C). These effects are much weaker than the ones observed when varying the other parameters, but since they have the same sign they will add up when we look at the total distance.

The s.d. *σ*_group distance_ of the two-dimensional Gaussian distribution for the distance between successive SFC groups has a twofold effect on the histogram distances (Fig 8D). Increasing *σ*_group distance_ increases the chance that later groups of an SFC leave the simulation volume. This reduces the chance of finding large patterns and the chance of finding long lags between pattern spikes, which is reflected in an increase of *d*_pattern size_ and dgrcup lags, respectively. However, dspatial distance decreases for larger *σ*_group_ distance. For a small s.d. there is a bias towards short spatial distances between pattern spikes, which is not what we observe in the experimental data. Since this parameter has opposite effects on multiple distances, we expect to find an optimal value which depends on the parameter set, and we expect the total distance to increase in both directions from that optimal value.

Additionally, we inspect how varying the number of SFCs embedded in the simulation volume *n*_chains_ influences how often single neurons participate in multiple SFCs (Fig 8E). We observe that *d*_multiple membership_ decreases from low to higher values of *n*_chains_, until it reaches a minimum at *n*_chains_ = 200. However, an even higher *n*_chains_ increases the chance for neurons to be involved in multiple patterns, since the number of chains is higher but the number of neurons stays the same. Neurons then have to be in more chains on average, which results in them being in more patterns, thus shifting the distribution of number of patterns a neuron appears in to the right (not shown). This measure thus constrains the total number of chains in our model, and we can use it to calculate the optimal number of embedded SFCs for each parameter set, which we will do in the following section. This will essentially allow us to eliminate one SFC parameter. We can do this since all other measures do not depend on *n*_chains_. For them, *n*_chains_ essentially determines the number of samples, but does not change the distribution. We see an increase towards the extreme case *n*_chains_ = 1 and some noise for very few chains, but then all measures except for dmultiple membership stabilize.

Finally, in (Fig 8F) we investigate the impact of the neuron density *ρ* on the histogram distances. *d*_pattern size_ increases slightly with increasing *ρ*, which matches our findings in Section 3.1.1, where we showed that increasing the total neuron density *ρ* decreases the SFC detectability. Since the number of chains and the number of neurons in a chain remains constant, but the total number of neurons increases, the detection probability decreases as the ratio of neurons in one chain to the total number of neurons goes down, thus reducing the chance to record two or more neurons from a chain. Here, this is reflected in an increase in *d*_pattern size_ since not enough large patterns are found to match the experimental data due to the decreased probability of recording multiple neurons from a chain. Besides this small effect on *d*_pattern size_, the neuron density does not show a significant impact on the histogram distances, since it does not affect the spatial distribution of SFC neurons in any way.

#### 3.2.3 Calculating the number of SFCs required to match the experimentally found spike pattern statistics

We have seen in the last section that the number of chains *n*_chains_ only affects the number of patterns a single neuron may participate in (*d*_multiple membership_). We saw a clear minimum of *d*_multiple membership_ at *n*_chains_ = 200 in Fig 8E. However, this optimal value also depends on *l* and *w*, since for a neuron to appear in the same number of SFCs, more chains are required if each chain contains fewer neurons (i.e. lower *l* or *w*). In order to verify that we can use dmultiple membership to find the optimal number of simulated SFCs *n*_chains_ for each parameter configuration, we plot *d*_multiple membership_ as a function of *n*_chains_ for all combinations of *l* and *w* in Fig 9. Varying the combination of *l* and *w* corresponds to varying the number of total neurons in an SFC which is simply the product *l* · *w*. We can see in Fig 9 that for every parameter combination, we find a clear minimum of *d*_multiple membership_ which we can use to fix *n*_chains_ to the corresponding number of simulated SFCs. The optimal values range from 40 for very long and wide chains containing many neurons each to 10, 000 for very narrow and short chains with few neurons per chain. In the following, we will fix *n*_chains_ for each parameter set using this method and only use the optimal value for all further plots and analyses.

**Fig 9.**
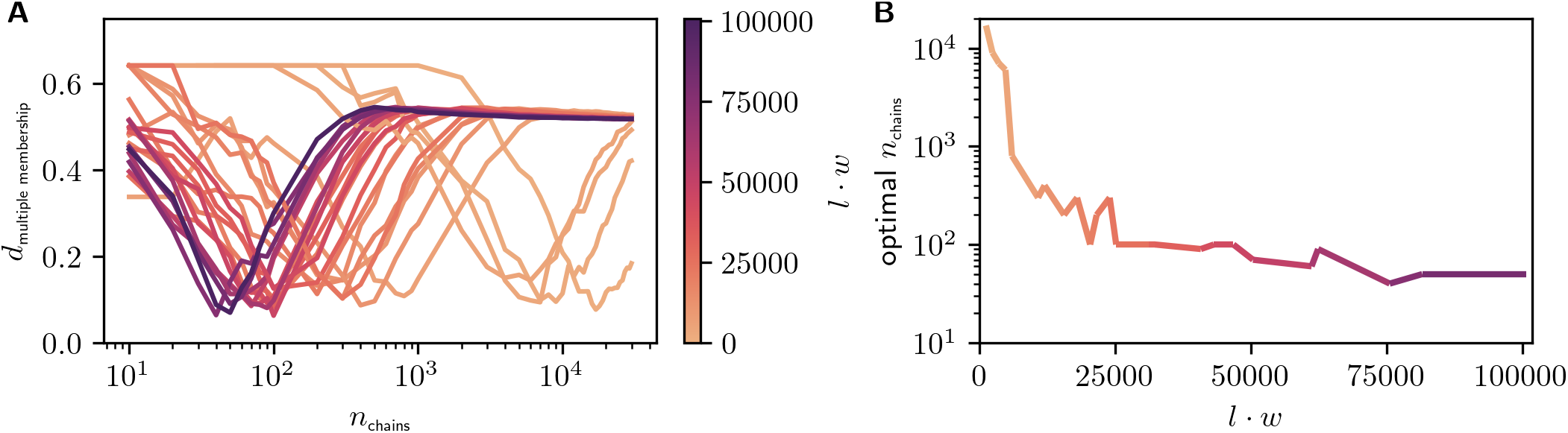
Fixing the number of SFCs by finding the minimal *d*_multiple membership_. (A) One line corresponds to a simulation run for fixed values of *l* and *w*. The product *l* · *w*, which corresponds to the number of neurons per SFC, is denoted by the line color. Along the x-axis, we vary the number of simulated SFCs *n*_chains_, and on the y-axis, the histogram distance for the multiple occurrences of neurons in SFCs is shown. Only data points for cases in which patterns have been detected are shown. No patterns were detected in the cases of chains with few neurons (low *l* · *w*) for very low values of *n*_chains_ ~ 10, which is very far from the optimal values of *n*_chains_ ~ 10^4^. (B) Optimal number of chains corresponding to the minimal *d*_multiple membership_ as a function of *l · w*, corresponding to the minima of the graphs in A.

We can estimate in how many chains a neuron participates in by dividing the number of “neuron slots” in all chains *l · w · n*_chains_ by the total number of neurons 840, 000 that exist in that cortical volume. Or in other words, single neurons have to participate in multiple chains to realize the chains. Thus, for the different parameter sets this estimate varies between 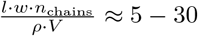 within the simulation volume *V* = 25.6 mm^3^ and the neuron density *ρ* = 35, 000 mm^-3^.

#### 3.2.4 Comparing model vs. experimental pattern statistics across the entire parameter space

Having fixed the optimal number of SFCs, and thus having eliminated one parameter, we can move on to a parameter scan to see how the total histogram distance behaves as a function of the remaining parameters. We showed in Section 3.2.2 that the neuron density does not have a significant impact on the histogram distances. Thus, we will focus on the SFC parameters *l, w, r*_group_, and *σ*_group distance_ and show the results in Fig 10.

**Fig 10.**
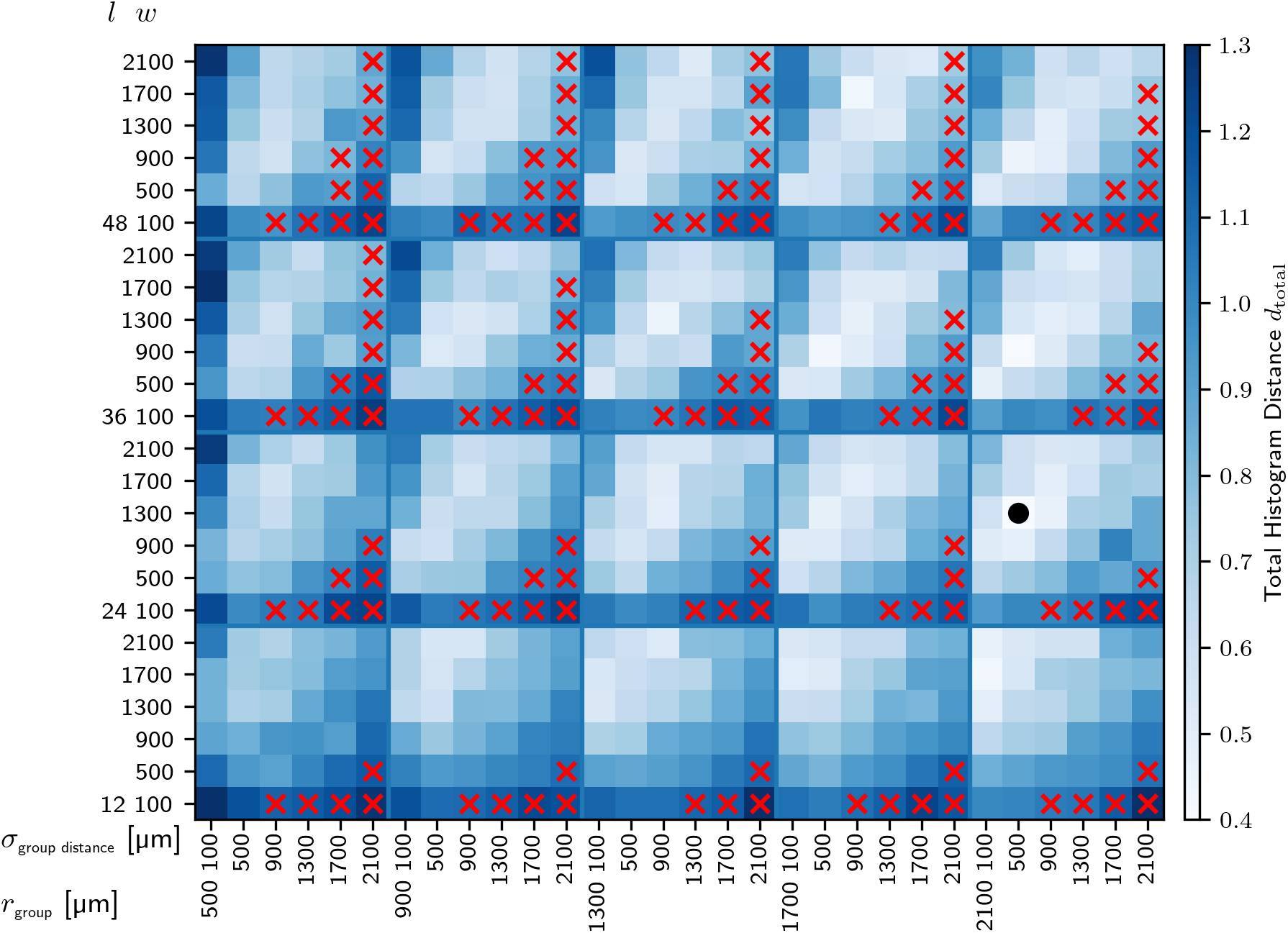
Total histogram distance between model and experimental results. Similar visualization to Fig 4. The color encodes the summed Euclidean distances between all histogram bins for the measures shown in Fig 11. Each entry corresponds to *N* = 20 runs of the simulated model. Red crosses indicate the parameter configurations where the chance of observing at least one pattern is lower than 80% (i.e., the lighter entries of Fig 4). The black dot represents the parameter configuration having the lowest total histogram distance.

Across the board, the histogram distance is quite consistently decreasing for increasing values of *r*_group_ (lighter shades of blue when moving box-by-box to the right). This is expected from Section 3.2.2, where we found that *r*_group_ has to be high to fit the distribution of time lags between pattern spikes.

In most sub-panels of Fig 10, for fixed *l* and *r*_group_, we see a diagonal band of lighter blue and even white shading, corresponding to smaller total histogram distances. This means that higher values of *σ*_group_ distance require higher values of *w* for a good fit. This effect has contributions from several single histogram distances, but can be seen most prominently for the pattern size histogram (see Fig 8). High values of *w* are required such that enough neurons per chain are detected to get large spike patterns as observed in the experimental data (see also Fig 8). However, there is a limit to this since at some point the patterns become larger than the experimental ones. This can be counter-balanced by increasing *σ*_group distance_, which decreases the pattern size since parts of the SFC are more likely to be outside of the simulation volume and are thus not detectable and cannot contribute to patterns. A higher SFC length *l* also increases the chance of larger patterns, which is why the diagonal of lighter shades moves down to lower values of *w* in the sub-panels for higher values of *l*.

As expected from Section 3.1.3, parameter sets with low *w* and high *σ*_group distance_ are ruled out since it is very unlikely that enough patterns can be detected in those cases. This corresponds to the parameter sets with a detection probability lower than 0.8 in Fig 7. These parameter configurations are indicated by red crosses in Fig 10. The corresponding total histogram distances are also rather high for these parameter sets as can be seen by the darker shades of blue in the respective sub-panels in the figure.

We have selected the parameter set with the lowest total histogram distance *d*_total_ = 0.40 (black dot) and we will look at its pattern statistics in the next section. This best fit of the experimental pattern statistics is reached at *l* = 24, *w* = 1, 300, *r*_group_ = 2,100 and *σ*_group distance_ = 500. As discussed in Section 3.2.2, a value of *l* > 12 does make sense here since it helps to increase the number of patterns with long temporal delays by increasing the number of possibilities to observe two neurons that are 12 SFC groups apart. In this parameter configuration with rather high values of *w* and *r*_group_, SFCs contain many neurons per group, and a single group covers most of the area of the Utah array. There is however a quite large region in the parameter space with an almost similarly good fit, so this is not the only possible setting in which the model can explain the experimentally found spike pattern statistics.

#### 3.2.5 Pattern statistics of the best-fit parameter set compared to the experiment

Finally, we compare the statistics of the patterns detected in the experiment to the statistics obtained in the best-fit parameter setting identified in the preceding section (Fig 11). Overall, the distribution of the best-fit parameter setting (in light blue) matches well with the experimental one (in orange) for most of the distributions (Fig 11A-C). The distribution of the Euclidean distance between pattern spikes (Fig 11D) is very variable due to the finite size of the electrode array. However, both the experimental and the model distribution show entries across most distances (see also Section 3.2).

**Fig 11.**
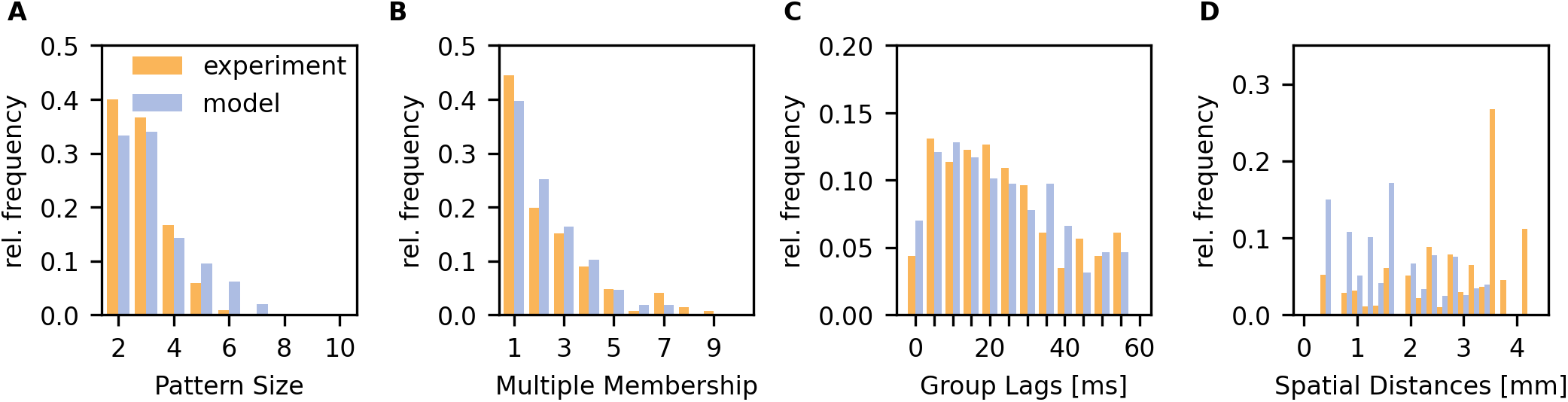
Pattern statistics in the experiment and the model. Experimental results are shown in orange, model results in light blue. The model parameters are *l* = 24, *w* = 1, 300, *r*_group_ = 2,100 and *σ*_group distance_ = 500. (A) Histogram of the number of pattern units (number of neurons involved in a pattern). (B) Histogram of number of patterns a single neuron appears in. Only neurons which take part in patterns are taken into account. (C) Histogram of time lags between neighboring pattern spikes. Only subsequent pattern spikes are taken into account. The temporal resolution of the histogram coincides with the one of the SPADE analysis, and is equal to 5 ms. The maximum time lag corresponds to the maximal pattern duration (here 60 ms). (D) Euclidean distance between pattern spikes. Only subsequent pattern spikes are taken into account. The histogram is normalized by the occurrences of the respective distances on the Utah array electrode grid.

## 4 Discussion

In this paper, we asked 1) whether synfire chains can be detected by Utah array recordings, and 2) whether or not the STPs and their statistics which we found in experimental data can be explained by synfire chains embedded below the Utah array. We were able to answer both questions positively, and found a large parameter range for the embedding of multiple SFCs that elicit the same STP statistics as the experimental data.

As data reference we used results on spatio-temporal spike patterns from 96-electrode recordings (Utah array) from monkey motor cortex (two monkeys) reported on in detail in a related study [40, 56]. The data contained between 80 and 152 simultaneously recorded neurons during motor behavior. Across 20 sessions (10 per monkey) we found STPs and summarized their statistics in four histograms with averaged data from all sessions of both monkeys (Fig 2): A) the number of neurons involved in STPs, B) the number of STPs a single neuron is involved in, C) the maximal temporal duration of STPs, and D) the spatial distances on the Utah array of the spikes involved in STPs.

We designed a model of the spatial embedding of one or more SFCs in the cortical area recorded from by the Utah array (motor cortex). The neuron composition of each SFC is parameterized by the length of the chain and the width of the groups. The distribution of the SFC groups in space is captured by the radius of a disk inside which all neurons of a single group are located, and by the standard deviation of a two-dimensional Gaussian distribution, which governs the positions of the group disk centers for subsequent groups. This choice of spatial distributions minimizes the model parameters. By varying these parameters we generated both localized and spatially extended SFCs in agreement with cortical connectivity.

Multiple anatomical and recording parameters enter in our calculations and introduce additional uncertainties. However, we were able to show in the appendix (S1 Appendix: The influence of the sensitive range of the electrodes) that the actual sensitive range of the electrodes does not have to be measured and is implicitly included in the number of isolated single units per electrode, a parameter available from experimental data, here 1.1. This is in agreement with [71—73] who report between 0.2 and 1.2 single units per electrode across multiple subjects and implantation sites. We define a model parameter *r*_sens_ to select neurons near the electrode tips which are candidates to be detected, but we were able to show that this parameter does not have any influence on the SFC detectability (Section 3.1.1). This is especially advantageous since the sensitive range has not been measured for Utah electrodes and thus underlies a large uncertainty. The neuron density *ρ* in the cortical layer in which the simulation volume is located constitutes another free parameter. However, we used the most conservative choice, i.e. the highest density (more neurons in total but same number of neurons within one SFC decreases the detectability, cf. Section 3.1.1), for our detectability calculation (cf. Section 3.1.3). Additionally, we showed that for the spike pattern statistics the density is not as critical since it does not affect the spatial distribution of SFC neurons (cf. Section 3.2.2).

We constrained the embedded SFCs to the cortical layer 2/3, which the experimental data was recorded from. Using the height of this layer as the height for the SFC volume, *h*_SFc Volume_ = 1.5 mm, (see [42–45]) and the neuronal density of *ρ* = 35,000 mm^-3^, we expect 840, 000 neurons inside this volume. In the experimental data we analyzed, 1.1 single units were detected per electrode, which we used as the basis for our neuron isolation model. This results in 110 recorded neurons, which corresponds to a subsampling of the neurons in our simulation of three orders of magnitude. We were able to show that despite this massive subsampling, SFCs can be detected in such a recording setting.

We distributed SFCs in the volume below the area of a Utah array and determined whether they can be detected by its electrodes. To this end, we evaluated the probability of recording single neurons, and we developed a detectability measure which requires at least two neurons from different groups of the same SFC to be recorded for the chain to be considered detected. Here, we assumed that every neuron of a chain always fires when the chain is activated, such that one SFC can only result in one STP

Going one step further, we predicted the statistics of the patterns that would be detected in such a setting and fitted the model embedding parameters to the pattern results obtained from the experimental data. We identified a region in the parameter space in which the spike pattern statistics that our model predicts match the experimental results well. The best match was found for a length of *l* = 24, a group size of *w* = 1, 300, a spatial radius of each group *r*_group_ = 2,100 μm and a group distance of *σ*_group distance_ = 500 μm, which corresponds to wide SFCs with many neurons per group and a broad spatial distribution. More combinations of *l*, *w*, *r*_group_ and *σ*_group distance_ provide comparable results, see Section 3.2.4.

Since more than one pattern was observed per session of experimental data and since some neurons took part in multiple patterns, we had to embed multiple SFCs at the same time and we had to allow neurons to be in more than one chain. We found an optimal match to the experimental data for broad spatial distributions since STPs in the experiment were detected across all possible spatial distances on the Utah array.

As shown in [74, 75], SFCs with large group sizes *w* and dense inter-group connectivity easily undergo an instability where even small fluctuations in the background activity trigger the formation of synchronous bursts of spiking activity. This problem of spontaneous synchronization in broad SFCs can be alleviated by a number of mechanisms, such as by diluting the inter-group connectivity [76], or by accounting for inhibitory feedback [18,77] or synaptic failure [78].

Despite the required large groups in our results, a dilution of inter-group connectivity to 100 to each receiving neuron in the following group guarantees that the activity runs stably through the chain ([76], and thus our assumption of stably running chains to find the amount of STPs as in the data would still be met.

Trengove et al. [18] were able to simulate networks in which 800,000 excitatory and 200,000 inhibitory neurons participate in a total of 51.020 SFC groups. Each neuron participates in ~ 70 chains. In their model, each SFC group activates a random set of inhibitory neurons, which regulates the network activity when a SFC is activated. Our scenario with 840,000 neurons participating on average in 5-30 SFC groups (see Section 3.2.3), depending on the parameter set, is close to the parameters of [18], and thus our setting appears to be feasible in dynamical simulations given that one implements a regulatory mechanism which is as effective as the inhibitory feedback employed in [18].

Since the electrodes of the Utah array have the same length, they likely recorded from neurons at the same cortical depth. So we have to assume that the observed phenomena are largely due to horizontal connectivity. The connection probability of two neurons mainly depends on the horizontal distance between their somata and drops off with increasing horizontal distance [79—83]. 80 — 82 % of the projection partners of a neuron in layer 2/3 commonly lie within a distance of 500 μm [79,84,85]. A previous SFC embedding study used a maximal distance of 300 μm between connected neurons as a constraint [76]. Our simulations that match the experimental data showed that large group radii (*r*_group_ ≥ 900 μm) and large inter-group distances (*σ*_group distance_ ≥ 500 μm) are required. As discussed above, such SFCs would have to be diluted, i.e. not fully connected, to function dynamically. This would help to stay within the anatomical bounds since a neuron does not have to be connected to every neuron of the next group and can instead just have connections to the closest neurons of the next group. Patchy connections of excitatory neurons in layer 2/3 cover ranges of 849.5 ± 337.5 μm [85—89], which would also account for the large group radii required in our model. However, a more recent study [84] showed in a dynamic photo-stimulation experiment lateral connections and high response reliability up to a range of 1,500 μm. These estimates cover the connection distances required for our optimal parameter set and thus constitute another candidate mechanism by which such chains could be realized.

Currently, we always fix all parameters to single values within a single simulation. However, a hybrid scenario of broad and narrow spatial profiles could be interesting and more realistic. SFCs with a broad spatial profile could serve information propagation within and between cortical areas, but in addition to this, there could be much more localized SFCs for local computations, which are connected to the grid of broader chains. Such a hybrid scenario, and interlinked chains in general, would increase the chance to find larger patterns. In [11,12], the authors found that information propagation along an SFC can be effectively gated by a second SFC, and, according to [16], the representation of information in spatially propagating structures is advantageous for information binding (cf. [14, 15]) across different cortical areas. These findings indicate that SFCs are a solid candidate for information propagation and computations on multiple scales, which would most likely require a multitude of spatial configurations of such interconnected chains. Recently, theoretical studies [21,22,26] have shown that an account of nonlinear synaptic integration in dendritic branches (dendritic action potentials) permits the formation of narrow SFC-like structures with only *w* ~ 10 neurons per group, supporting a reliable propagation of small volleys of synchronous spikes, and thereby constitutes an efficient mechanism for learning and processing of complex sequences of data.

In the same data set that we analyzed for STPs, [90] find a spatial structure of correlated neurons (measured by covariance) which changes across behavioral contexts. The covariances are partly strong within neurons up to few mm distance. This goes well with the fact that we found STPs involving neurons across all possible distances on the 4 × 4 mm^2^ Utah array. However, it is currently not clear if neurons involved in an active SFC also increase their rates. Grün (private communication, based on data from [91]) showed that neurons involved in an SFC seem to go hand in hand with coherent rate changes as visible in the population raster, however, when neurons are sorted according to their spike times in the SFC, it turns out that these are actually single well timed spikes.

A limitation of the SPADE analysis lays in the fact that the method is able to detect only spike patterns repeating exactly (within few ms imprecision) across the data. Thus, it does not allow for temporal imprecision in the spike sequences over a few milliseconds, and does not allow for selective participation of the pattern neurons. In other words, single pattern instances, where some spikes may be missing due to synaptic failure, due to e.g. imprecision in the spike sorting, are not detected by SPADE. However, other approaches relying on different methodologies may detect such “fuzzy patterns” [92–96]. Thus, the results obtained by the analysis of the experimental data may be compared to analysis of other pattern detection methods, perhaps leading to more patterns and/or to somewhat different statistics.

In order to keep the model as simple as possible, we did not include the significance testing performed in SPADE [51,54] either. Modeling it more closely could also improve the match of the pattern size distributions, however, it is almost impossible without fully dynamical simulations, since the rates of the neurons and thus the numbers of occurrences of the patterns are important factors required for the significance test.

Our model can also be adapted to different recording settings. The electrode positions and sensitive ranges, depth of the recorded cortical layer and the local neuron density however have to be provided. Using other recording techniques and applying our analyses could help to answer remaining and follow-up questions. Neuropixels probes provide a very good spatial resolution orthogonal to the surface of the cortex. Data recorded with such probes could be analyzed and used to model SFCs across multiple cortical layers. A high sampling resolution along the other dimensions is however much more difficult to achieve with such probes since it would require multiple probes to be implanted close to one another.

STPs have been detected in many studies across different animals, cortical areas, temporal scales and task conditions. Riehle et al. [29] found synchronous patterns at 330 μm to 660 μm distance that would correspond to neurons recorded from a single SFC group. The implied group radius is in accordance with many parameter sets that provide a good fit to the data we analyzed, too. They also found synchronous activations related to behavior of different subsets within a set of three neurons, hinting at multiple participation of neurons in SFCs which we also assume to fit our data. In the same data we analyzed, Torre et al. [31] also found synchronous firing of neurons at distances of 400 μm to 2, 400 μm, implying similar group radii. Prut et al. [28] found spatio-temporal spike patterns with different maximum temporal extents ranging from 0 ms to 500 ms within a cortical area. Modeling the latter with SFCs would require very long chains or much larger delays between groups. They even found different inter-spike delays between spikes of the same neurons in different patterns, adding to the evidence for multiple participation of neurons in SFCs. Hemberger et al. [32] found fuzzy spiking sequences which could be triggered by single spikes. The fact that a single spike from one neuron suffices to trigger a reliable spiking sequence also hints at the possibility of very narrow chains with few neurons per group and with strong synapses, which can still be reliable if a single spike is already enough to propagate the activity to the following groups.

As follow-ups to this work, SFC spatial embeddings should be complemented by including spiking activity to study the dynamics. A closer modeling of pattern significance testing may also help to match the experimentally found pattern size distribution even better. A possible outlook is to apply our model to more spike recordings where evidences of STP activity has been found, to investigate whether different parameter values are required to match pattern statistics in other experimental settings.

## Code and Data Availability Statement

The code and data to perform and reproduce the analyses presented in this study can be found at https://dx.doi.org/10.5281/zenodo.6840944.

## Acknowledgments

We thank Markus Diesmann for valuable discussions.

## Supplementary material

### S1 Appendix: The influence of the sensitive range of the electrodes

In our three-dimensional model, the isolation probability of a neuron depends on the sensitive range *r*_sens_ of the electrode and the neuron density *ρ* (see Eq 3):

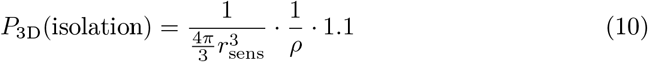

Note that the “real” sensitive range is implicitly included in the experimentally obtained figure of 1.1^single units/electrode^ (cf. Section 2.1). Our parameter *r*_sens_ is required in the model to select the neurons close to electrode tips which are candidates to be detected. Our isolation probability *P*_3D_ (isolation) is the number of single units per electrode divided by the total number of neurons within the sensitive range of the electrode.

This isolation probability applies to neurons which are inside the sensitive range of an electrode. So, to get the total number of isolated neurons, we have to multiply this probability by the number of neurons which are inside the sensitive ranges of electrodes. Assuming a homogeneous neuron density, this number exactly corresponds to the denominator in Eq 10:

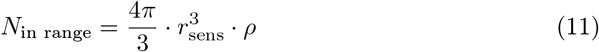

Thus, the total number of isolated neurons is independent of our parameter rsens:

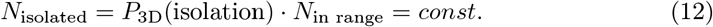

This does not mean that the actual sensitive range of the electrodes does not matter, it is hidden in the 1.1^single units/electrode^ in Eq 10. Our model parameter *r*_sens_ is not equal to the actual sensitive range, and it does not have to be, since via Eq 10 we ensure that the overall isolation probability of neurons per electrode in our model matches the experimental data. In order to perform the simulations, we have to assign a value to *r*_sens_ in order to be able to select neurons close to electrode tips, and we fix *r*_sens_ = 50 μm.

### S2 Figure: Impact of SFCs leaving the simulation volume

Scanning *σ*_group_ as in Fig 5, once as before (black dots indicating the mean with orange area indicating the standard error of the mean) and once with periodic boundary conditions (gray dots indicating the mean with turquoise band indicating the standard error of the mean). See Fig 5 and description thereof for an explanation of the simulation procedure. In the case with periodic boundary conditions, SFCs that leave the simulation volume enter it again on the opposite side.

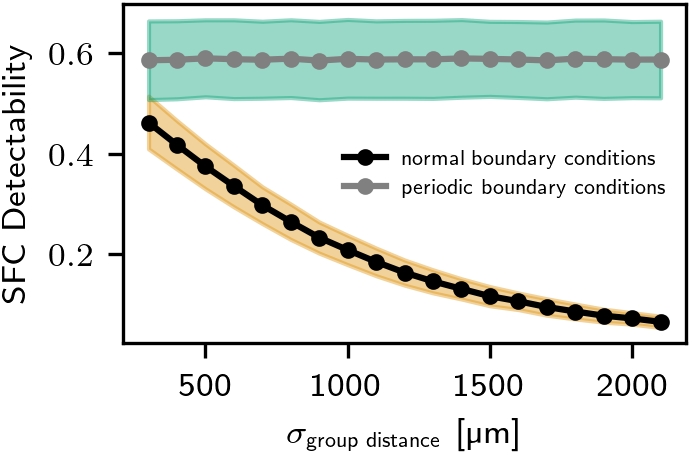

